# Transcriptome analysis uncovers a link between copper metabolism, and both fungal fitness and antifungal sensitivity in the opportunistic yeast *Candida albicans*

**DOI:** 10.1101/2020.03.27.011775

**Authors:** Inès Khemiri, Faiza Tebbji, Adnane Sellam

## Abstract

Copper homeostasis is an important determinant for virulence of many human pathogenic fungi such as the highly prevalent yeast *Candida albicans*. However, beyond the copper transporter Ctr1, little is known regarding other genes and biological processes that are affected by copper. To gain insight into the cellular processes that are modulated by copper abundance in *C. albicans*, we monitored the global gene expression dynamic under both copper depletion and excess using RNA-seq. Beyond copper metabolism, other different transcriptional programs related to fungal fitness such as stress responses, antifungal sensitivity, host invasion and commensalism were modulated in response to copper variations. We have also investigated the transcriptome of the mutant of the copper utilization regulator, *mac1*, and identified potential direct targets of this transcription factor under copper starvation. We also showed that Mac1 was required for the invasion and adhesion to host cells and antifungal tolerance. This study provides a framework for future studies to examine the link between copper metabolism and essential functions that modulate fungal virulence and fitness inside the host.

## Introduction

The redox properties of copper (Cu) make this trace element crucial for biological systems as it serves as an essential cofactor of enzymes that functions in many biological processes including iron acquisition, antioxidative defense and energy metabolism (Festa and Thiele, 2011). Cu is also toxic for the cell and its accumulation should be tightly monitored and kept at homeostatic levels. In the context of host-pathogen interaction, Cu is thought to be elemental for what is known as nutritional immunity of host cells during fungal infections (Hood and Skaar, 2012; Samanovic et al., 2012; Djoko et al., 2015). Cu is differentially distributed across different anatomical sites of the human body where it is mostly abundant in skeleton and bone marrow, skeletal muscle, liver, brain and blood (Linder et al., 1998). Host cells seems to use both Cu-sequestration and Cu-poisoning to limit fungal pathogens from proliferating in different niches. In the case of the meningitis causing agent *Cryptococcus neoformans*, Cu is available at limiting concentrations in the brain interstitium while it is highly abundant in lung airways (Ballou and Wilson, 2016). For the highly prevalent human pathogenic yeast *Candida albicans*, Cu availability is highly dynamic for the same anatomical niche within the host. For instance, upon earlier phase of kidney colonization, *C. albicans* is confronted by high levels of Cu which is considered as a host-imposed Cu-poisoning strategy (Mackie et al., 2016; Culbertson et al., 2020). This earlier Cu spike is followed by a rapid sequestration by renal tissues to probably limit fungal growth as *C. albicans* activates its own Cu utilization machinery to promote its fitness.

Our current knowledge on regulatory mechanisms of Cu homeostasis of eukaryotic cells came mainly from works in the model yeast *Saccharomyces cerevisiae*. When Cu becomes limiting, *S. cerevisiae* promotes Cu import through the activation of the transcription factor Mac1 that induces the transcription of the high affinity Cu membrane transporters Ctr1 and Ctr3 (González et al., 2008). Transcript levels of the ferric reductase Fre1 and Fre7 that reduces Cu (Cu^2+^➔ Cu^+^) prior to its uptake by Ctr1 is also activated by Mac1 (van Bakel et al., 2005). Response to Cu excess is primarily controlled by the transcription factor Cup2 that activates different Cu-chaperones such as Cup1-1, Crs5 and Ccc2 in addition to the Cu-transporting P-type ATPase that are required for Cu tolerance (González et al., 2008).

Pathogenic fungi are exposed to a labile pool of Cu within the human host and has consequently evolved a tight regulatory control to ensure cellular homeostasis of this trace element. As in *S. cerevisiae*, *C. albicans* uses Mac1 to mediate the activation of both the Cu transporter Ctr1 and the ferric reductase Fre7 under Cu starvation (Marvin et al., 2003; Woodacre et al., 2008). Furthermore, *C. albicans* uses the P-type ATPase Crp1 that function as a Cu extrusion pump to survive in high Cu environments (Weissman et al., 2000). *C. albicans* has an ortholog of Cup2 that is required for Cu tolerance (Homann et al., 2009), however, its role as a transcriptional modulator of Cu detoxification has not been explored so far. Interestingly, under Cu excess, *C. albicans* activates the Cu-dependent superoxide dismutase Sod1 (Cu-Sod1) to neutralize the superoxide anion while it uses the Mn-requiring Sod3 (Mn-Sod3) under Cu limitation (Li et al., 2015). As Sod enzymes use metals as cofactors to convert superoxide to oxygen and hydrogen peroxide, *C. albicans* shifts metal co-factors for superoxide dismutase depending on Cu abundance in the colonized niches. *C. neoformans* depends on the Cu-transporters Ctr1 and Ctr4 for Cu uptake and on the Cu-metallothioneins MT1 and MT2 for Cu detoxification (Ding et al., 2011, 99). In this pathogenic fungus, both Cu uptake and detoxification are governed by the same transcriptional regulator Cuf1 (Ding et al., 2011, 99; Garcia-Santamarina et al., 2018). In *Aspergillus fumigatus*, intracellular Cu uptake relies on the Cu-transporters *ctrA2* and *ctrC* that are both under the control of the transcriptional factor *Afmac1* (Cai et al., 2017). Alteration of either uptake or detoxification processes in *C. albicans*, *A. fumigatus*, *C. neoformans* and the dimorphic fungus *Histoplasma capsulatum* impairs fungal virulence and fitness (Waterman et al., 2007; Ding et al., 2013; Sun et al., 2014; Mackie et al., 2016; Cai et al., 2017) suggesting that Cu metabolism might be a promising therapeutic targeted to treat fungal infections.

While Cu homeostasis is an important virulence determinant in *C. albicans* (Mackie et al., 2016), the impact of Cu availability on the transcriptome of this important human pathogen remain unexplored. Furthermore, beyond the Cu transporter Ctr1, little is known regarding other genes or biological process that are affected by Cu abundance or modulated by Mac1 at the genome level in *C. albicans.* To gain insight into the cellular processes that are modulated by Cu abundance in *C. albicans*, we monitored the global gene expression dynamic under both Cu depletion and excess using RNA-seq. Our data uncovered that, in addition to Cu utilization genes, other different cellular processes related to fungal fitness were modulated in response to Cu fluctuations. We have also investigated the transcriptome of *mac1* mutant and identified potential direct targets of this transcription factor under Cu starvation. We also showed that Mac1 was required for the invasion and the adhesion to human enterocytes and antifungal tolerance. Thus, this study provides a framework for future studies to examine the link between Cu metabolism and essential functions that modulate fungal virulence and fitness inside the host.

## Materials and Methods

### Fungal strains and media

*C. albicans* was routinely maintained at 30°C on YPD (1 % yeast extract, 2 % peptone, 2 % dextrose, with 50 mg/ml uridine). The *C. albicans* WT strain SN250 (*ura3*/::imm434/*URA3 iro1*::*IRO1*/*iro1*::imm434 *his1*::*hisG*/*his*::*hisG leu2*::Cd*HIS1*/*leu2*::Cm*LEU2 arg4*/*arg4*) (Noble et al., 2010) used in this study derives from the SC5314 clinical strain. *mac1* (*mac1*::*LEU2*/*mac1*::*HIS1*) deletion mutant come from the transcription factor deletion collection (Homann et al., 2009) that was constructed in SN125 (*his1*/*his1*, *leu2*/*leu2*, *arg4*/*arg4*, *URA3*/*ura3*::imm434, *IRO1*/*iro1*::imm434).

To build the *mac1* complemented strain, the *MAC1* gene was reintegrated into the null mutant *mac1* strain using pDUP3 plasmid (Gerami-Nejad et al., 2013). Briefly, the *MAC1* locus containing its endogenous promoter ([-678,0] intergenic region) was amplified by PCR using primers containing flanking sequences homologous to pDUP3. The resulting pDUP3-*MAC1* construct was digested by *Sfi*I and integrated into the *NEU5L* genomic site of the *mac1* strain as previously described (Gerami-Nejad et al., 2013) using lithium acetate transformation (Wilson et al., 2000). Transformants were selected on YPD plates supplemented with nourseothricin at 200 μg/ml nourseothricin and correct integration was verified by PCR. Primers used for *MAC1* cloning in pDUP3 plasmid and for the diagnosis of pDUP3-*MAC1* integration are listed in the **Supplementary Table S1**.

### Growth inhibition assays

All chemicals used in this study were provided by Sigma-Aldrich (St. Louis, MO, USA). Working stock solutions of Copper (II) Sulfate (CuSO_4_; 532 mM), Bathocuproinedisulfonic acid disodium salt (BCS; 8 mM), Bathophenanthrolinedisulfonic acid (BPS; 8 mM) and fluconazole (3.3 mM) were prepared using Milli-Q sterile water. Amphotericin B and miconazole stock solutions were prepared using dimethyl sulfoxide (DMSO; Sigma-Aldrich) at a concentration of 1 mM and 2 mM, respectively.

For growth assays in liquid YPD medium, overnight cultures of *C. albicans* were resuspended in fresh YPD medium at an OD_600_ of 0.05 and added to a flat-bottom 96-well plate in a total volume of 100 μl per well in addition of the tested compounds. For each experiment, a compound-free positive growth control and a cell-free negative control were included. Growth assay curves were performed in triplicate in 96-well plates using a Sunrise plate-reader (Tecan) at 30°C under constant agitation with OD_600_ readings taken every 10 min for 48h.

For spot dilution assays, overnight cultures were diluted to an OD_600_ of 0.1 and fivefold serial dilutions were prepared in distilled water. A total of 4 μl of each dilution were spotted on YP-agar with different carbon sources (glucose, maltose, glycerol and ethanol) at 2 % or on YPD plates with the different antifungals: amphotericin B (1 μg/ml), miconazole (0.5 μg/ml) and fluconazole (0.5 μg/ml). Plates were incubated at 30°C for 3 days and imaged using the SP-imager system.

### Expression analysis by RNA-seq and qPCR

Overnight cultures of *mac1* mutant and WT (SN250) strains were diluted to an OD_600_ of 0.1 in 40 ml of fresh YPD-uridine medium and grown at 30°C under agitation (200 rpm) to an OD_600_ of 0.4. Cultures were then either left untreated or exposed to either CuSO_4_ (2mM) or BCS (400μM), and incubated at 30°C for 30 min. For each condition, a total of two biological replicates were considered for RNA-seq analysis. Cells were then harvested by centrifugation at 3,000 x g for 5 min and the pellets were quick-frozen and stored at −80°C. Total RNA was extracted using an RNAeasy purification kit (Qiagen) and glass bead lysis in a Biospec Mini 24 bead-beater. Total RNA was eluted, assessed for integrity on an Agilent 4200 TapeStation System prior to cDNA library preparation. The NEBNext® UltraTM II RNA Library Prep Kit for Illumina was used to construct the RNA-seq library following the manufacturer’s instruction. The quality, quantity and the size distribution of the libraries were determined using an Agilent Bioanalyzer. A 2×100 paired-end sequencing of cDNAs were performed using an Illumina Novaseq6000 sequencing system.

For qPCR confirmation experiments, a total of three biological and three assay replicates were performed. cDNA was synthesized from 1μg of total RNA using High-Capacity cDNA Reverse Transcription kit (Applied Biosystems). The mixture was incubated at 25°C for 10 min, 37°C for 120 min and 85°C for 5 min. 2U/μl of RNAse H (NEB) was added to remove RNA and samples were incubated at 37°C for 20 min. qPCR was performed using a LightCycler 480 Instrument (Roche Life Science) for 40 amplification cycles with the PowerUp™ SYBR® Green master mix (Applied Biosystems). The reactions were incubated at 50°C for 2 min, 95°C for 2min and cycled for 40 times at 95°C, 15 s; 54°C, 30 s; 72°C, 1 min. Fold-enrichment of each tested transcripts was estimated using the comparative ΔΔCt method. To evaluate the gene expression level, the results were normalized using Ct values obtained from Actin (*ACT1*, C1_13700W_A). Primer sequences used for this analysis are summarized in **Supplementary Table S1.**

### RNA-sequencing data analysis

Adaptor sequences and low quality score bases (Phred score < 30) were first trimmed using Trimmomatic (Bolger et al., 2014). The resulting reads were aligned to the *C. albicans* reference assembly SC5314 from Ensembl (http://fungi.ensembl.org/Candida_albicans_sc5314_gca_000182965/Info/Index), using STAR (Dobin et al., 2013). Read counts are obtained using HTSeq (Anders et al., 2015) and are represented as a table which reports, for each sample (columns), the number of reads mapped to a given gene (rows). For all downstream analyses, we excluded lowly-expressed genes with an average read count lower than 1 across all samples, resulting in 5,942 genes in total. The R package *limma* (Ritchie et al., 2015) was used to identify differences in gene expression levels between treated and non-treated samples. Nominal *p*-values were corrected for multiple testing using the Benjamini-Hochberg method. Differentially expressed transcripts in the **Supplementary Tables S2** and **S4** were identified using a false-discovery rate (FDR) of 5% and 2-fold enrichment cut-off. Gene ontology (GO) analysis was performed using GO Term Finder of the Candida Genome Database (Skrzypek et al., 2017). The GSEA Pre-Ranked tool (http://www.broadinstitute.org/gsea/) (Subramanian et al., 2005) was used to determine statistical significance of correlations between the characterized transcriptomes with a ranked gene list or GO biological process terms as described by Sellam *et al*. (Sellam et al., 2014a).

### HT-29 adherence and damage assay

Damage to the human colon epithelial cell line HT-29 was assessed using a lactate dehydrogenase (LDH) cytotoxicity detection kit^PLUS^ (Roche), which measures the release of the LDH enzyme in the growth medium. HT-29 cells were grown in 96-well plate as monolayers in McCoy’s medium supplemented with 10 % FBS at 1×10^4^ cells per well and incubated at 37°C with 5 % CO_2_ overnight. HT-29 cells were then infected with *C. albicans* cells at MOI cell:yeast of 1:2 in the presence or the absence of BCS (100 μM or 800 μM), for 24 h at 37°C with 5 % CO_2_. HT-29 cells were also incubated with 100 μM or 800 μM BCS alone to rule out any toxic effect of this compound on HT-29 cells. No discernable growth defect of HT-29 cells was noticed with both BCS concentrations. Following incubation, 100 μl of supernatant was removed from each experimental well and LDH activity in this supernatant was determined by measuring the absorbance at 490 nm (OD_490_) following the manufacturer’s instructions.

For the adherence assay, *C. albicans* cells were co-incubated with HT-29 cells at 37°C and 5 % CO_2_ for 1 hour. Non-adherent cells were removed by rinsing five times with 1 ml PBS and cells were then fixed with 4 % paraformaldehyde. HT-29 cells were permeabilized with 0.5 % Triton X-100 and adherent fungal cells were stained with 2 μM calcofluor white during 30 min in the dark at room temperature. Adherent cells were visualised using Cytation 5 high-content microscope with 20x magnification and DAPI filter. For each well, at least 10 fields were photographed.

## Results

### Global transcriptional responses to copper depletion and repletion in *C. albicans*

To gain insight into the cellular processes that are modulated by Cu abundance in the pathogenic yeast *C. albicans*, we monitored the global gene expression dynamic under both Cu depletion (400 μM BCS) and excess (2mM CuSO_4_) using RNA-seq. Transcripts associated with both Cu utilization and detoxification were identified by comparing the transcriptional profiles of WT cells treated with BCS and CuSO_4_, respectively, to that of non-treated WT cells. In response to Cu excess, *C. albicans* activated transcripts associated with Cu detoxification and transport including the cytosolic small chaperone *ATX1*, the metallothionein *CUP1*, the P-type ATPase *CCC2* and the Cu efflux pump *CRP1* (**Figure 1A-B**). We also found that genes of Cu uptake such as the Cu transporter *CTR1*, the transcription factor *MAC1* and the cupric reductase *FRE7* were downregulated (**Figure 1A**). Transcripts involved in iron uptake, and both cytosolic and mitochondrial ribosome biogenesis were activated whereas those related to glycolysis, energy and, NAD and hexose/glucose metabolisms were repressed (**Figure 1B**). Genes related to drug response and detoxification such as the MFS transporter *MDR1*, as well as its transcriptional activator *MRR1*, together with other MFS (*FLU1*, *NAG3*, *NAG4*, *TPO3*, *TPO4*), ABC (*MLT1*) and MaTE (*ERC1*, *ERC3*) transporters were significantly induced. Among repressed transcripts, we also found many genes of the ergosterol biosynthesis pathway (*ERG1*, *ERG3*, *ERG11*, *ARE2*) in addition to their *bona fide* transcriptional regulator Upc2 (**Supplementary Table S2**).

**Figure 1.**
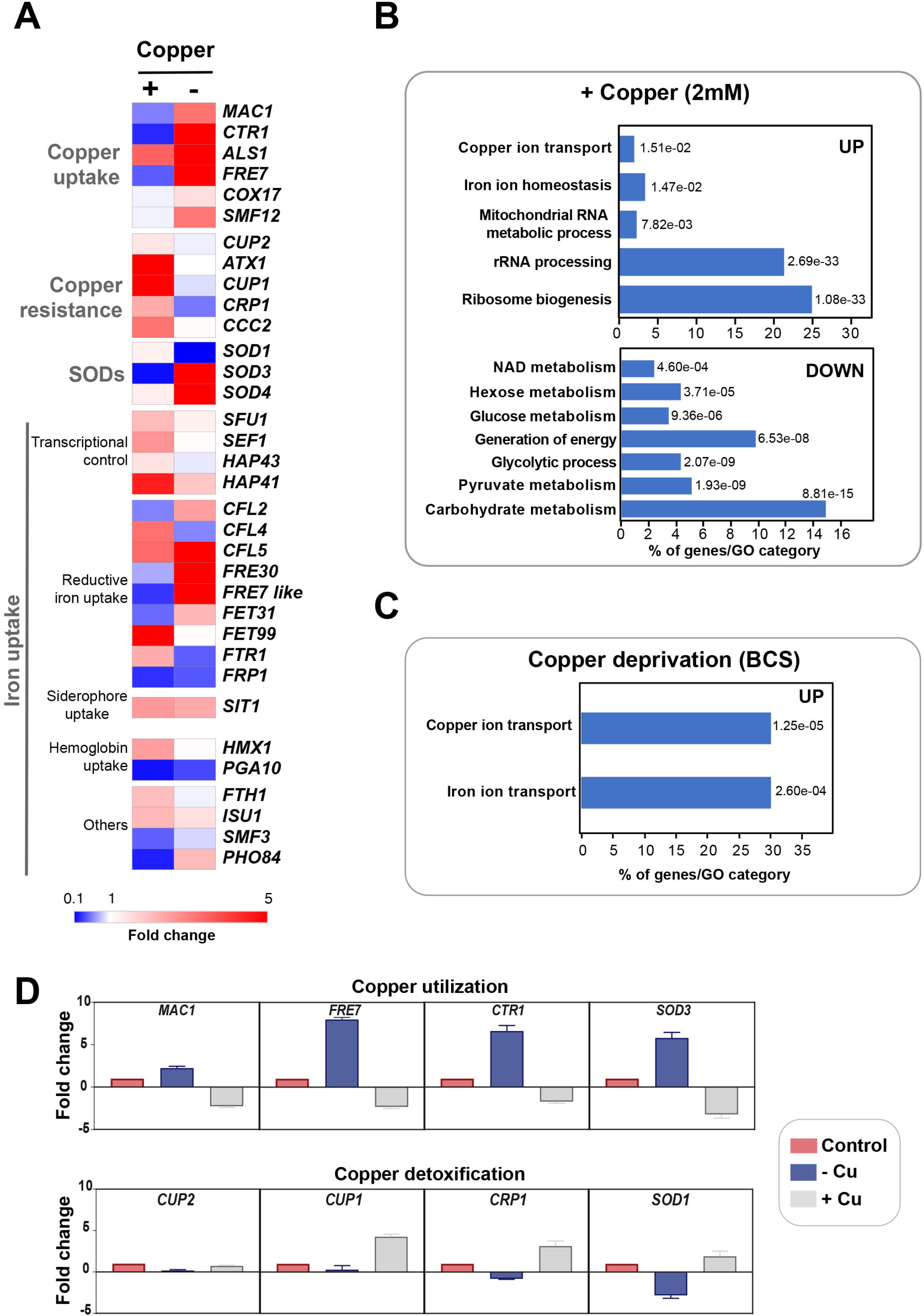
Genome-wide transcriptional profiling of *C. albicans* to Cu variations by RNA-seq. (**A**) Heat map visualization of the transcript levels of the Cu homeostasis pathway in *C. albicans* in response to Cu excess and deprivation. *C. albicans* WT cells were exposed to either 2 mM CuSO_4_ or 400 μM BCS, and incubated at 30°C for 30 min. Transcripts associated with both Cu utilization (copper “−”) and detoxification (copper “+”) were identified by comparing the transcriptional profiles of WT cells treated with BCS and CuSO4, respectively, to that of non-treated WT cells. (**B-C**) Gene function and biological process enriched in the transcriptional profiles of *C. albicans* growing under Cu excess (**B**) and limitation (**C**). (**D**) qPCR validation of RNA-seq data. Transcript levels of both Cu utilization (*MAC1*, *CTR1*, *FRE7* and *SOD3*) and detoxification (*CUP1*, *CRP1*, *SOD1* and *CUP2*) genes were assessed and fold-changes were calculated using the comparative ΔΔCt method. Data were normalized using Ct values obtained from actin gene in each condition.

Under Cu deprivation, as in other fungi, *C. albicans* upregulated genes required for Cu internalization including the Cu transporter Ctr1 and the ferric reductases Fre7, Fre30 and orf19.7077 that reduce Cu to facilitate its uptake by Ctr1 (**Figure 1A, 1C** and **Supplementary Table S2**). Other transcripts including the manganese transporter *SMF12* and the ABC transporter *SNQ2* were also activated. We also found that under Cu depletion, *C. albicans* upregulated the Mn-dependant superoxide dismutase *SOD3* (Mn-*SOD3*) and repressed the Cu-dependant *SOD1* (Cu-*SOD1*). Reversely, under Cu excess, Cu-*SOD1* were activated and Mn-*SOD3* repressed. This switch from Cu to Mn cofactors of Sods is an adaptative strategy for *C. albicans* cells to resist to reactive oxygen species (ROS) inside the host when Cu become limiting (Li et al., 2015). Taken together, our data suggest that Cu abundance did not affect exclusively Cu detoxification and utilization pathways but also other biological processes that might require this essential micronutrient to fulfill their functions.

qPCR confirmed gene expression alterations as shown by RNA-seq for both Cu utilization (*MAC1, CTR1, FRE7* and *SOD3*) and detoxification (*CUP1*, *CRP1*, *SOD1* and *CUP2*) genes under both Cu deprivation and excess, respectively (**Figure 1D**).

### Gene set enrichment analysis of the *C. albicans* Cu-sensitive transcriptomes

Gene Set Enrichment Analysis (GSEA) was used to mine the transcriptomes associated with both Cu starvation and excess, to uncover potential resemblance with *C. albicans* genome annotations and other experimental large-scale omics data (Subramanian et al., 2005; Sellam et al., 2014b) (**Supplementary Table S3**). **Figure 2A** and **2C** summarize the significant correlations of the Cu-responsive transcriptomes with either functional GO categories, *C. albicans* transcriptional signatures in different physiological conditions or in specific mutant background, and also with genome-wide promoter occupancies data. For both Cu availability conditions, GSEA recapitulated most of the GO correlations as identified by the conventional methodology. Cells growing in Cu excess exhibited a transcriptional profile similar to that of different mutants of transcription factors such as *upc2*, *efg1* and *ace2* and to mutants of different signaling pathways including Ras1-cAMP-PKA (*ras1*) and the AGC protein kinase Sch9 pathways in addition to a significant correlation with a set of genes with protomers enriched in Cat8 binding motif (**Figure 2A-B** and **Supplementary Table S3**).

**Figure 2.**
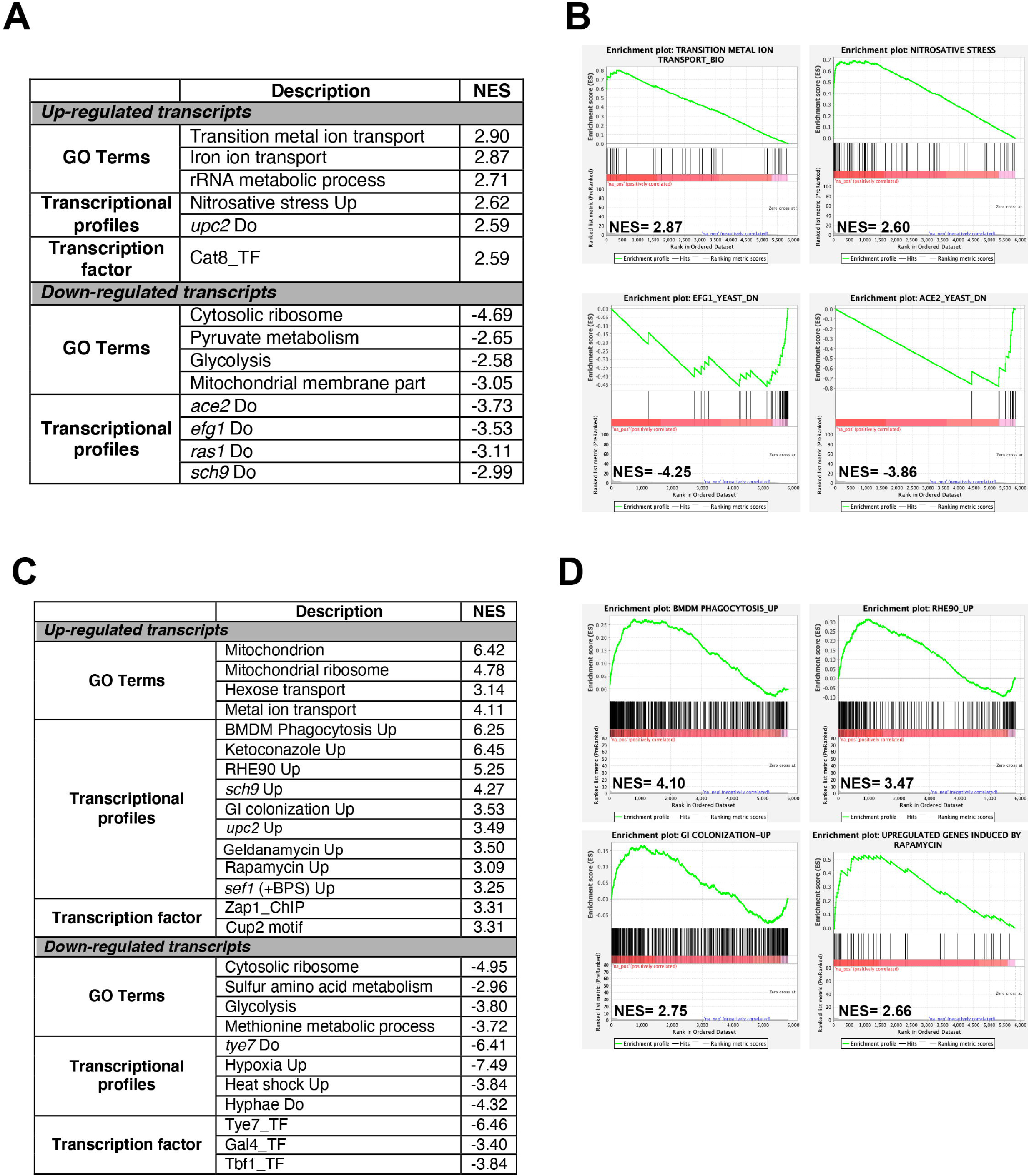
Gene set enrichment analysis of RNA-seq data. RNA-seq data of Cu excess (**A**-**B**) and deprivation (**C**-**D**) were analyzed using Gene Set Enrichment Analysis (GSEA). Relevant correlations between *C. albicans* transcriptome under Cu excess (**A**) and limitation (**C**) and other gene sets and functions are summarized. The complete GSEA correlations are listed in **Supplementary Table S3**. Graphs of GSEA for both Cu abundance conditions were also shown (**B** and **D**). NES, normalized enrichment score.

Under Cu deprivation, upregulated transcripts were significantly similar to the *C. albicans* transcriptional programs expressed during the colonization of the murine gut (Pierce et al., 2013), and the interaction with host cells including the human oral epithelial cells (Spiering et al., 2010) and the bone marrow-derived mouse macrophages (Marcil et al., 2008). This similarity suggests that, during the interaction with the host, *C. albicans* might be confronted by a Cu-deprived environment that host might generate as an immune strategy to sequester this essential metal. Transcriptional similarities were also observed with the transcriptomes associated with fungal fitness and virulence traits including persistence under hypoxia and biofilm formation or during exposures to antifungal molecules (ketoconazole, rapamycin and geldanamycin) (**Figure 2C-D** and **Supplementary Table S3**). As in other human pathogenic fungi, *C. albicans* might connect the control of fitness and virulence attributes to the Cu cellular adaptive machinery as a cue to promote either commensalism or pathogenicity (Ballou and Wilson, 2016).

### Mac1-mediated control of copper deprivation

Prior to assessing the contribution of the transcription factor Mac1 to the global response of *C. albicans* to Cu deprivation, we first tested and confirmed the growth defect of *mac1* mutant under different conditions including Cu deprivation, utilization of non-fermentable carbon sources and iron chelation as previously reported (Marvin et al., 2004, 1; Homann et al., 2009; Khamooshi et al., 2014) (**Figure 3**). To identify Mac1-dependant transcripts associated with Cu utilization, we compared the transcriptional profile of *mac1* cells exposed to BCS to that of non-treated *mac1* cells. Overall, our RNA-seq data showed that the inducibility of genes of Cu uptake and utilization including *CTR1*, *ALS1*, *FRE17*, *COX17* in addition to the Mn-*SOD3* and many iron uptake transcripts (*CFL2*, *CFL5*, *FRE30*, *FET31* and orf19.7077) were lost in *mac1* mutant under Cu depleted conditions (**Figure 4A**). This data confirmed the role of Mac1 as the *bona fide* transcriptional regulator of Cu homeostasis in *C. albicans*.

**Figure 3.**
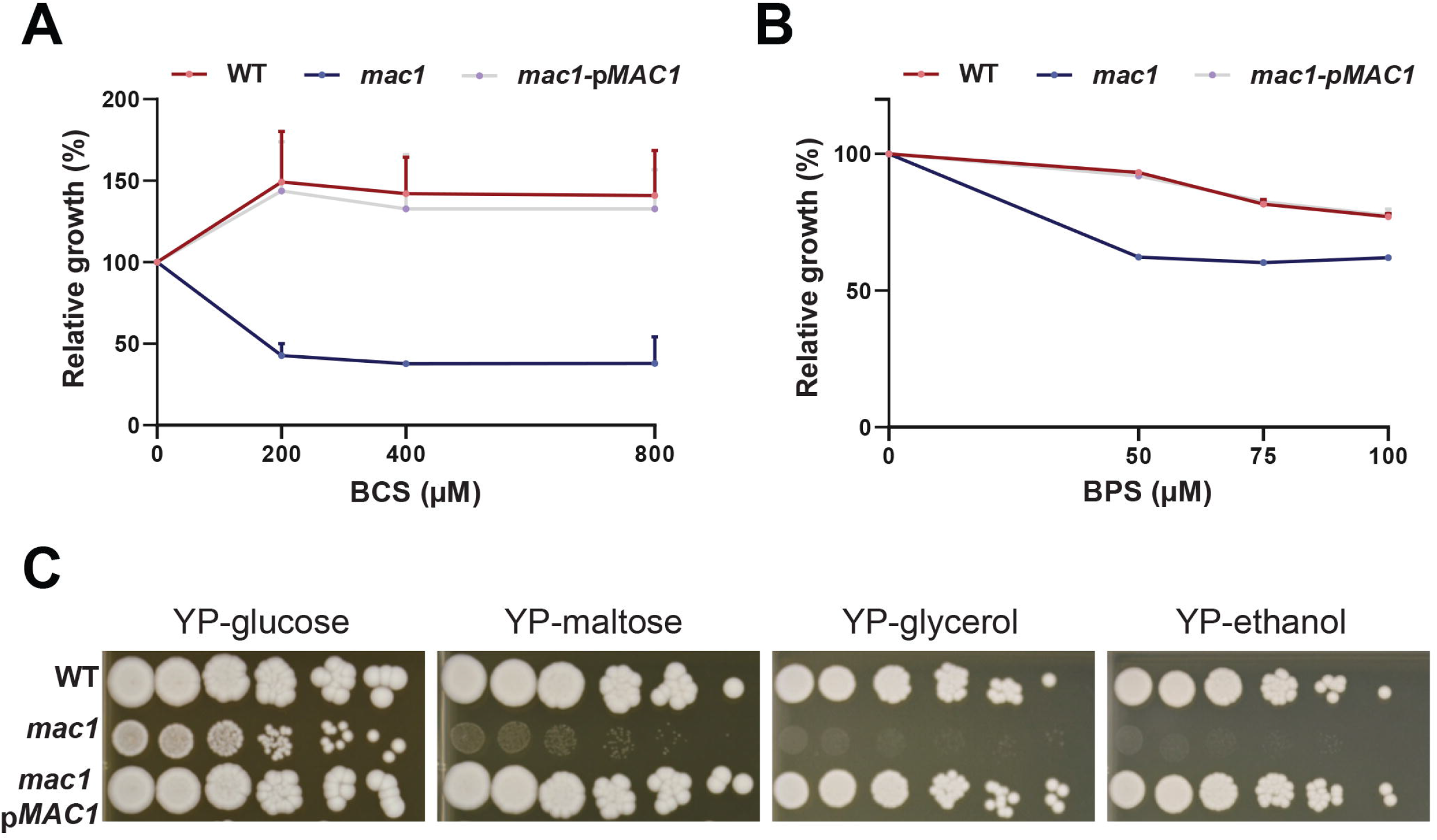
Phenotypic characterisation of *mac1* mutant. (**A-B**) The *C. albicans* WT, *mac1* and the *mac1* complemented strains were grown at 30°C in YPD supplemented with different concentrations of BCS or BPS. OD_600nm_ reading was taken after 48 h of incubation. ODs measurements for each concentration of BCS and BPS are the mean of triplicate. (**C**) *mac1* is required for the utilization of alternative and non-fermentable carbon sources. The *C. albicans* WT, *mac1* and the *mac1* complemented strains were serially diluted, spotted on media with the indicated carbon sources and incubated for 3 days at 30 LJC.

**Figure 4.**
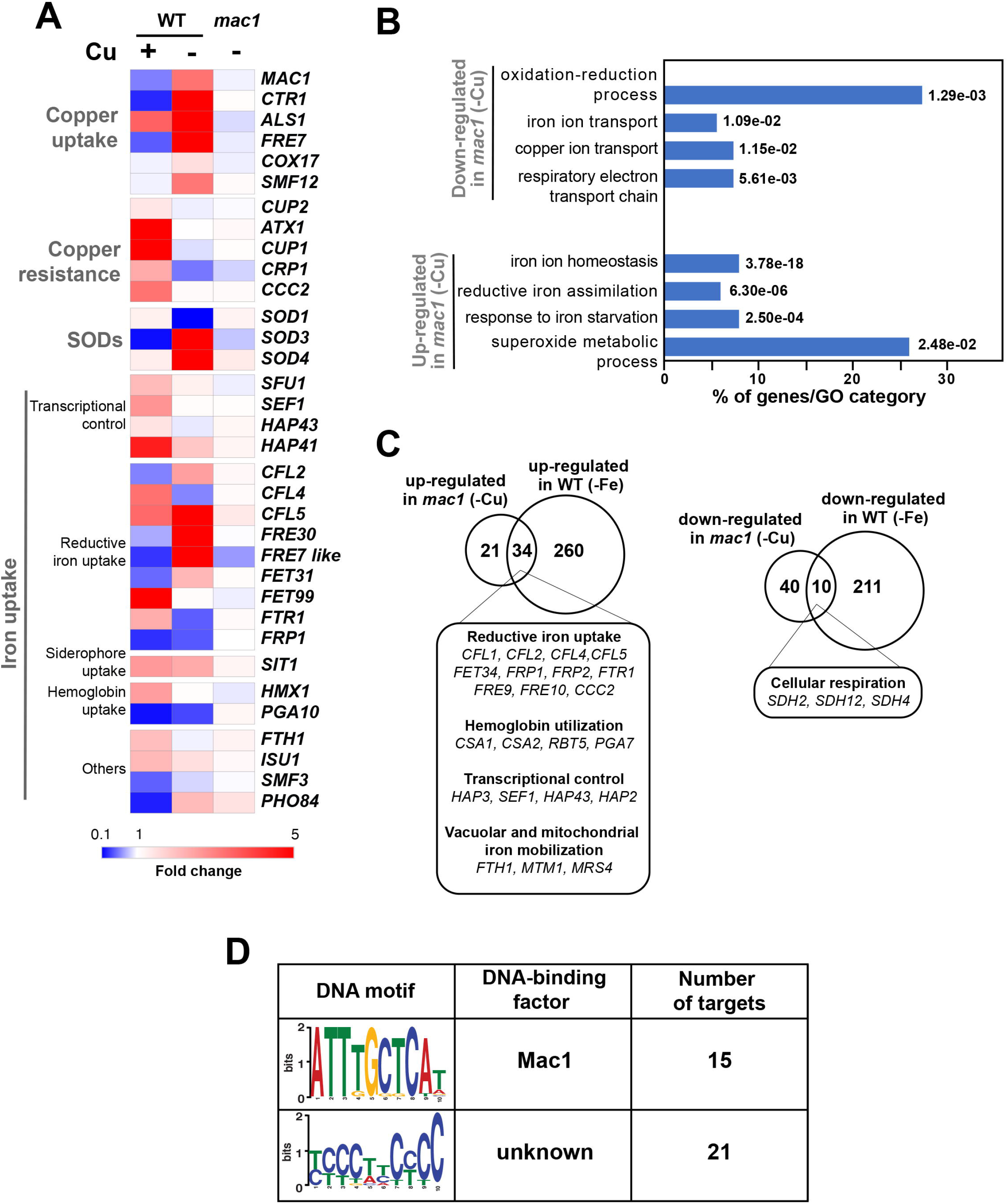
The Mac1-dependant control of Cu deprivation transcriptome. (**A**) Transcriptional alterations of Cu homeostasis genes in both WT and *mac1*. Heatmap visualisation was presented as in **Figure 1A** for the WT and includes *mac1* RNA-seq data. As the WT strain, *mac1* mutant were exposed to 400 μM BCS, and incubated at 30°C for 30 min. Mac1-dependant transcripts associated with Cu utilization were identified by comparing the transcriptional profile of *mac1* cells exposed to BCS to that of non-treated *mac1* cells. (**B**) Gene ontology analysis of upregulated and downregulated transcripts in *mac1* in response to BSC. The *p*-values were calculated using hypergeometric distribution. (**C**) Comparison of the Cu- and the iron-depletion transcriptomes of *C. albicans* (from Chen *et al.* 2011). Venn diagrams summarize the similarity between up- and downregulated transcripts under both iron and Cu starvations. Functional gene categories enriched in both conditions are indicated. (**D**) *De novo* prediction of DNA cis-regulatory motif enriched in the 50 gene promoters that *mac1* failed to activate using MEME analysis tool (http://meme-suite.org).

Other cellular processes were also altered in *mac1* mutant as compared to the WT under Cu deprivation (**Figure 4B** and **Supplementary Table S4**) including the repression of respiratory electron transport chain genes and the upregulation of iron homeostasis processes (**Figure 4A-C**). This transcriptional signature is reminiscent of an iron deprived environment that is most likely due to the fact that many iron uptake and utilization proteins depend on Cu for their functionality (Knight et al., 2002). Accordingly, a total of 34 among the 55 upregulated transcripts in *mac1* were also upregulated in *C. albicans* cells under iron starvation as previously shown (Chen et al., 2011) and most of those genes are under the control of the Sfu1-Sef1-Hap43 iron transcriptional control axis (**Figure 4C**). Of note, we found that the transcriptional level of Sef1 and the CCAAT-binding factors Hap3, Hap1 and Hap43 were activated in *mac1* mutant.

We also analyzed the 50 gene promoters that *mac1* failed to activate for potential consensus binding motifs and identified two enriched mini-sequences one of which is the well-known Mac1 binding site (McDaniels et al., 1999; Garcia-Santamarina et al., 2018) (**Figure 4D**). Mac1 binding motif was found in the promoters of Ctr1 and Sod3, as previously reported (Woodacre et al., 2008; Li et al., 2015), in addition to the ferric reductases Fre30 and Fre7 (**Table 1**). Mac1-regulatory motif was also found in 5’ cis-regulatory regions of genes that are not related directly to Cu metabolism such as electron transport chain (Sdh2, Sdh4, Cyb2), nitrosative stress response (Yhb1) and, manganese (Smf12) and iron transport (Ftr1) genes.

**Table 1.** List of genes with putative Mac1 cis-regulatory motif.

| Gene name | Orf19 ID | Motif location |  | Description |
| --- | --- | --- | --- | --- |
|  |  | Start | End |  |
| *CTR1* | orf19.3646 | -274 | -266 | copper transporter |
| *FTR2* | orf19.7231 | -501 | -493 | high-affinity iron permease |
| *FRE30* | orf19.6140 | -237 | -229 | protein with similarity to ferric reductases |
|  | orf19.7078 | -893 | -885 | ortholog of *S. cerevisiae* YCL012C, proteins localize to the endoplasmic reticulum and vacuole |
| *FRE7 like* | orf19.7077 | -133 | -125 | putative ferric reductase |
| *CWH43* | orf19.3225 | -261 | -253 | putative sensor/transporter protein with a predicted role in cell wall biogenesis |
| *TIM12* | orf19.4620 | -202 | -194 | component of the Tim22 complex, involved in protein import into mitochondrial inner membrane |
|  | orf19.1409.2 | -668 | -660 | protein of unknown function with predicted transmembrane domain |
|  |  | -949 | -941 |  |
| *FDH1* | orf19.638 | -243 | -235 | formate dehydrogenase; oxidizes formate to CO<sub>2</sub> |
| *YHB1* | orf19.3707 | -834 | -824 | nitric oxide dioxygenase; acts in nitric oxide scavenging/detoxification |
|  | orf19.6277 | -423 | -415 | protein of unknown function, has a PhoX domain with phosphoinositide-binding activity |
| *SDH4* | orf19.4468 | -265 | -257 | succinate dehydrogenase |
| *SMF12* | orf19.2270 | -279 | -271 | manganese transporter |
| *CYB2* | orf19.5000 | -138 | -130 | putative cytochrome b2 precursor |
| *SOD3* | orf19.7111.1 | -153 | -145 | cytosolic manganese-containing superoxide dismutase |
| *SDH2* | orf19.637 | -297 | -289 | succinate dehydrogenase, Fe-S subunit |

### Copper modulates antifungal sensitivity and virulence

GSEA analysis of the *C. albicans* transcripts modulated by Cu limitation uncovered a significant correlation with the transcriptional program expressed when cells are challenged with the azole antifungal, ketoconazole (**Figure 2C**). This led to hypothesize that both stresses affect similar cellular processes in *C. albicans* or that Cu homeostasis modulates drug sensitivity. We found that genetic inactivation of *MAC1* rendered *C. albicans* cells hypersensitive to both polyene (amphotericin B) and azole (fluconazole and miconazole) antifungals (**Figure 5A**). Antifungal sensitivity of *mac1* was reversed by supplementing the growth medium by CuSO_4_ which suggests that Cu homeostasis modulates antifungal tolerance in *C. albicans* (**Figure 5B**).

**Figure 5.**
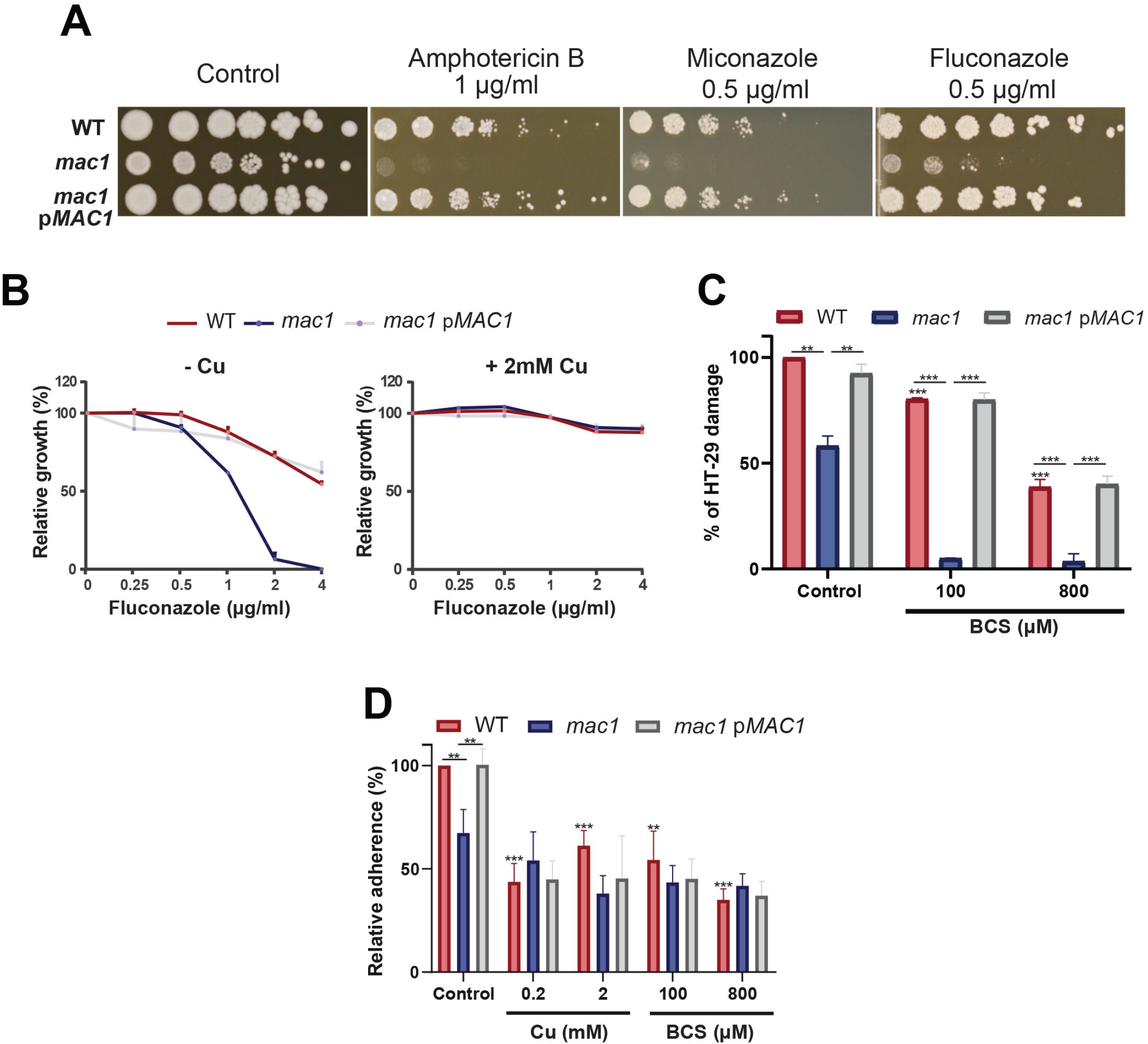
Copper modulates antifungal sensitivity and host invasion. (**A**) Mac1 modulates antifungal sensitivity. *C. albicans* WT, *mac1* and *mac1* complemented strains were serially diluted, spotted on media with different antifungals (amphotericin B, miconazole and fluconazole) and incubated for 3 days at 30 ◻C. (**B**) Cu supplementation restore *mac1* growth inhibition by fluconazole. The *C. albicans* WT, *mac1* and the *mac1* complemented strains were grown at 30°C on YPD with different concentrations of fluconazole with or without 2 mM CuSO_4_. OD_600_ reading was taken after 48 h of incubation. (**C**) *MAC1* inactivation and Cu depletion attenuate damage of the human colon epithelial HT-29 cells. HT-29 cell damage was assessed using the lactate dehydrogenase (LDH) release assay and was calculated as percentage of LDH activity in cell infected by *mac1* and the revertant strains, with or without BCS, relatively to cells infected by the WT (SN250) strain. At least four biological replicates were obtained for each experiment. (**D**) *mac1* mutant had a reduced adherence to HT-29 cells. *C. albicans* WT, *mac1* and *mac1* complemented strains were co-incubated with HT-29 for one hour at 37°C and 5 % CO_2_. Fungal cells were stained with 2 μM calcofluor white and counted. Data are presented as means ± SD from at least three independent experiments performed in triplicate. Statistical difference for each conditions vs. WT control or between conditions was determined by two-tailed Student’s t test. **P<0.01, ***P<0.005.

As the transcriptional profile of *C. albicans* cells under Cu deprivation is similar to that experienced during the interaction with the human host (**Figure 2C-D**), we tested whether Mac1 is required to damage the HT-29 human enterocytes using the LDH release assay. *mac1* mutant exhibited a reduced ability to damage HT-29 cells as compared to the WT and the revertant strains (**Figure 5C**). HT-29 damage by *mac1* mutant was further attenuated when both *C. albicans* and host cells were cocultured in the presence of BCS. Interestingly, Cu chelation exerts a protective antifungal activity since supplementation of the culture medium with 100 and 800 μM of BCS prevented 20 and 62 % of HT-29 damage by the *C. albicans* WT strain, respectively. Together these data suggest that Cu homeostasis in *C. albicans* and the availability of this metal in host niches is essential for fungal virulence.

The adherence of *mac1* mutant to the HT-29 enterocytes was also tested and the obtained data indicate a significant reduction of *mac1* attachment as compared to the WT and the complemented strain (**Figure 5D**). Moreover, adding or depriving Cu from the growth medium reduced significantly the WT adherence to the HT-29. *mac1* adherence defect to HT-29 was not reverted by Cu supplementation or phenocopied by Cu chelation in the WT strain (**Figure 5D**). This suggests that Mac1 might control other biological processes independently from its *bona fide* role as Cu metabolism modulator.

## Discussion

So far, the global response of *C. albicans* to Cu availability remain unexplored. Previous works have focused only on known Cu metabolic genes that are homologous to *S. cerevisiae* such as the Cu transporter Ctr1 (Marvin et al., 2004, 1), the transcription factor Mac1 (Woodacre et al., 2008) and the Cu efflux pump Crp1 (Weissman et al., 2000). Here, we explored the genome-wide transcriptional response of *C. albicans* to elevated Cu or Cu deprivation in order to unbiasedly assess cellular processes associated with fungal fitness that are modulated by Cu availability.

Our RNA-seq data recapitulated the *bona fide* fungal response to Cu. When grown under limiting concentrations, *C. albicans* activate Cu utilization genes such as the Cu transporter Ctr1 while under Cu excess genes involved in Cu detoxification (Crp1) and relocalization (Atx1, Cup1, Ccc2) were induced. As in other fungi (van Bakel et al., 2005; Rustici et al., 2007; Garcia-Santamarina et al., 2018), we also found that iron utilization genes were differentially modulated by Cu depletion which might reflect a situation of iron deficiency as well. Indeed, iron uptake depends on the multicopper ferroxidases and consequently Cu deprivation engender a subordinate iron depletion in *C. albicans* cells. Intriguingly, upon Cu excess, transcript levels of the transcription factor Sef1 and its direct target genes related to iron uptake and utilization such as the high affinity iron permease Ftr1, the ferric reductase Cfl5, the siderophore transporter Sit1 and the heme oxygenase Hmx1 were upregulated. This transcriptional signature reflects a defect in iron internalization as was also noticed under Cu limitation. This is similar to what was observed in yeast and mammalian cells where Cu overload deplete intracellular iron levels (Arredondo et al., 2004; Jo et al., 2008). This phenomenon could be explained by the fact that, as eukaryotic cells try to detoxify Cu by efflux, this essential metal become limiting for the Cu-dependent iron uptake machinery which led to iron deficiency.

The main goal of the current study was to uncover, beyond the Cu homeostatic routes, other biological processes that are modulated by Cu. Mining the *C. albicans* Cu transcriptome uncovered significant similarities with transcriptional programs activated by this opportunistic yeast in different niches within the host or during the interaction with immune cells. This highlight the importance of Cu for both fungal fitness and survival to immune response. Accordingly, our data showed that depriving *C. albicans* cells from Cu or inactivating the master transcriptional regulator of Cu uptake *MAC1* both led to decreased damage and adherence to human enterocytes. Virulence defect in these conditions could be attributed to the fact that Cu is important for *C. albicans* to express its virulence factors such as metabolic flexibility and stress resistance. As shown previously (Marvin et al., 2004, 1; Mackie et al., 2016) and confirmed by our work, a homeostatic level of Cu is essential for *C. albicans* to metabolize different carbon sources, an important attribute for this yeast to colonize diverse niches with contrasting nutrients (Miramon and Lorenz, 2017; Burgain et al., 2019). Furthermore, as Sod1 requires Cu for the disproportionation of superoxide into peroxide and oxygen, Cu depletion might deteriorate the ability of *C. albicans* to resist the oxidative burst killing by phagocytic immune cells (Kang et al., 2002; Frohner et al., 2009).

At the genome level, our data uncovered that *C. albicans* cells grown in BCS exhibited a transcriptional pattern similar to that of cells challenged with ketoconazole. This suggest that Cu starvation, as azoles, might lead to sterol depletion. In accordance with that, we have shown that the Cu deficient mutant *mac1* was hypersensitive to the ergosterol inhibitors (miconazole, fluconazole and amphotericin B) as did sterol deficient mutants such as *upc2* (Silver et al., 2004; MacPherson et al., 2005; Hoot et al., 2011). The opposite is also possible where sterol decrease might lead to Cu deprivation. However, a recent study had shown that ergosterol depletion by fluconazole do not affect the intracellular amount of Cu in *C. albicans* (Hunsaker and Franz, 2019). The growth defect of *mac1* mutant in the presence of fluconazole was reverted by Cu supplementation suggesting that Cu might correct ergosterol depletion by promoting *de novo* synthesis of this fungal sterol. In a support of such hypothesis, studies in *S. cerevisiae* has shown that Cu promote ergosterol biosynthesis by promoting the transcription of *ERG* genes which led to the rescue of growth inhibition by lovastatin, an inhibitor of the ergosterol biosynthetic enzyme Hmg1 (Fowler et al., 2011).

This current investigation provides a rich framework for future work to uncover novel candidates for Cu homeostasis. For instance, under Cu excess, different categories of transporters were induced suggesting their contribution to Cu efflux. Under Cu starvation, the manganese transporter *SMF12* were induced and it is a potential target of Mac1 as its promoter has a putative Mac1-binding motif. Thus, in addition to manganese, Smf12 might serve as a Cu transporter as was suggested also in *S. cerevisiae* (Liu et al., 1997). Furthermore, our data indicate that in addition to the *bona fide* Cu uptake genes, Mac1 might have other potential new targets and modulate other biological functions (**Table 1**) such as the succinate dehydrogenase complex activity (*SDH12*, *SDH2*, *SDH4*), mitochondrial protein translocation and insertion in inner membrane (*TIM12*) and adhesion (*ALS1*). Why Mac1 might link Cu metabolism to those functions remains to be determined. Taken together, these assumptions should provide a fertile area for future investigations.

## Author Contributions

IK, FT and AS contributed to the design, execution, and/or data collection, and analysis of the experiments of this work. AS drafted the manuscript. All authors contributed to revision and final approval of the manuscript.

## Supplementary Material

The datasets generated for this study are available at Figshare: https://figshare.com/s/98f0075055b2737ad1f5

## Supplementary data

**Supplementary Table S1.** List of primers used in this study.

**Supplementary Table S2.** Transcripts differentially expressed in *C. albicans* WT (SN250) under Cu excess (+Cu) and limitation (+BCS) using a 1.5-fold change cut-off and a 5% false discovery rate.

**Supplementary Table S3.** Gene Set Enrichment Analysis (GSEA) of the *C. albicans* WT (SN250) transcriptome under Cu excess (+Cu) and depletion (+BCS).

**Supplementary Table S4.** Transcripts differentially expressed in *mac1* under Cu depletion (+BCS) using a 1.5-fold change cut-off and a 5% false discovery rate.

**Supplementary Table S5.** RNA-seq raw data of both *C. albicans* WT strain (under Cu excess and limitation) and *mac1* mutant (under Cu limitation).

